# Nuclear inhibitor of protein phosphatase 1 (NIPP1) regulates CNS tau phosphorylation and myelination during development

**DOI:** 10.1101/2022.04.02.486849

**Authors:** Cody McKee, Peter Shrager, Arindam Mazumder, Archan Ganguly, Abigail Mayer, Karl Foley, Nancy Ward, Margaret Youngman, Hailong Hou, Houhui Xia

**Author notes:** **Corresponding author**: Dr. Houhui Xia, Dr. Hailong Hou.

## Abstract

Nuclear inhibitor of protein phosphatase 1 (NIPP1) is a known regulator of gene expression and has been shown to play roles in many physiological or pathological processes such as stem cell proliferation and skin inflammation. While NIPP1 has many regulatory roles in proliferating cells, its function in the central nervous system (CNS) has not been directly investigated. In the present study, we examined NIPP1 CNS function using a conditional knockout (cKO) mouse model, in which the *nipp1* gene is excised from neural precursor cells. These mice demonstrate severe developmental impairments that lead to premature lethality within the first few postnatal weeks. To delineate some of the neurological changes occurring in these animals, we first assessed microtubule associated protein tau, a known target of NIPP1 activity. Furthermore, observed tremors prompted exploration of myelin integrity, an integral structure for CNS function, whose disruption is associated with various neurological disorders and neurodegenerative diseases. First, immunoblotting demonstrated increased phospho-tau and altered AKT and PP1 activity in NIPP1 cKO mice, suggesting increased tau phosphorylation likely results from a shift in kinase/phosphatase activity. Second, immunoblots, electron microscopy, and electrophysiology demonstrated a myelin deficit within the brain and optic nerve. Our study suggests that NIPP1 in neural precursors regulates phosphorylation of tau and CNS myelination and may represent a novel therapeutic target for neurodegenerative diseases.

## Introduction

Nuclear inhibitor of protein phosphatase 1 (NIPP1) is a known regulator of protein phosphatase 1 (PP1) targeting PP1 to sub-nuclear localization [1]. Despite its name, PP1 in NIPP1 complex is active in vivo [2]. Recently NIPP1 has been shown to regulate gene expression and can interact directly with epigenetic machinery, specifically EZH2 of the polycomb repressive complex 2 (PRC2) on chromatin [3–6]. For PRC2-target genes, NIPP1 likely functions as a gene suppressor. However, many genes, whose expression is regulated by NIPP1, are not PRC2-target genes [6], suggesting that NIPP1 can also regulate gene expression independently of PRC2. NIPP1 plays important roles in many physiological or pathological processes throughout the body, such as early embryonic development/cell proliferation [7], progenitor cell expansion in the adult liver [8], mammalian spermatogenesis [9], and chemokine-driven skin inflammation [10]. Recent studies suggest a role for NIPP1 in the regulation of neuronal proteins including tau [11, 12], implying a previously unexplored role for NIPP1 within the CNS.

A critical protein linked to NIPP1 activity and neurodegenerative diseases is microtubule associated protein, tau. Tau has many functions including stabilizing microtubules and facilitating fast axonal transport [13]. Tau phosphorylation is an active and crucial area of research due to the abundance of hyperphosphorylated tau in many neurodegenerative diseases including Alzheimer’s disease (AD) and frontotemporal dementia with parkinsonism-17 (FTDP-17) [13]. The phosphorylation status of tau is a balance between kinases and phosphatases, including protein phosphatase-1 [14]. Whether nuclear inhibitor of protein phosphatase-1 (NIPP1) regulates tau phosphorylation is not yet clear.

In this report, we investigated the function of NIPP1 within the CNS by deleting NIPP1 in embryonic neural progenitor cells (NPCs), under the nestin promoter. We find that Nestin-Cre;*nipp1*^fl/fl^ (cKO) mice exhibit substantial developmental impairments, and a tremor similar to that described in animals with disrupted myelination [15]. Western blots probed for tau demonstrate a major increase in tau phosphorylation as well as alterations in tau associated phosphatase/kinase activation, including PP1 and AKT. Moreover, due to the presence of a tremor and poor coordination we explored myelin structure and function in different regions of the CNS. We find that NIPP1 cKO mice show clear deficits in myelination within the brain and optic nerve as assayed by western blots, electron microscopy and electrophysiology. Thus far, our work demonstrates an important CNS function for NIPP1 in tau hyperphosphorylation and myelination, possibly by regulating gene expression and phosphorylation signaling pathways.

## Results

We studied the potential function of NIPP1 in the CNS by deleting the *nipp1* gene in NPCs by crossing a Nestin-Cre mouse line (Jackson Laboratories) with our floxed *nipp1* mouse line [8]. The resulting Nestin-Cre;*nipp1*^fl/fl^ mouse (NIPP1 cKO) leads to embryonic knockout of NIPP1 protein in neurons, astrocytes, and oligodendrocytes. The mouse pups are visibly smaller than their wild type (WT) littermates (Fig. 1A) and typically die prior to postnatal day (P) 30. Western blotting experiments confirmed almost complete NIPP1 protein knockout in the cortex, hippocampus and many other brain regions (Fig. 1B). These data also demonstrate that any expression by non-NPCs is minimal.

**Figure 1.**
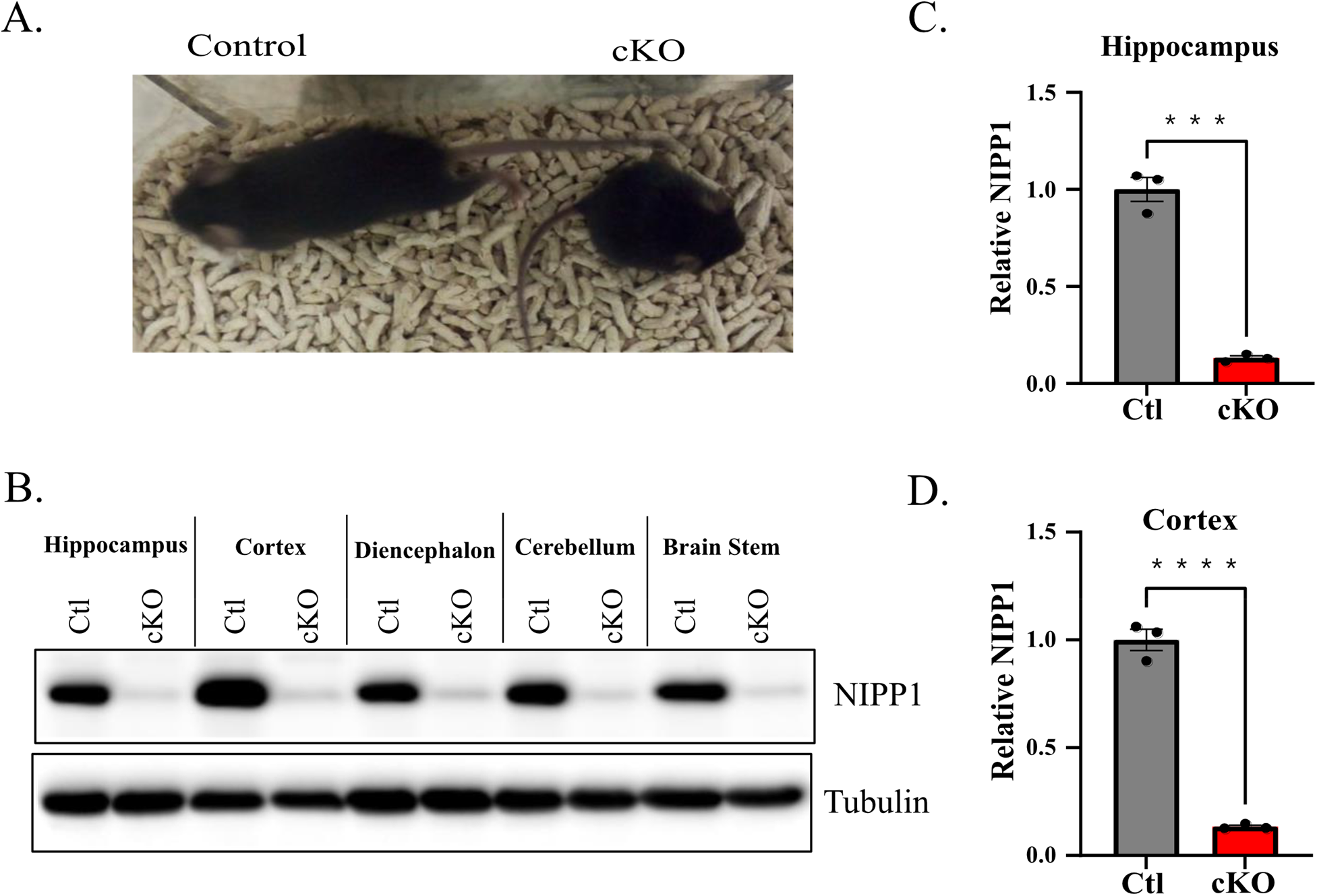
NIPP1 cKO mice are smaller than littermate controls. **A.** Image of control mouse (left) with NIPP1 cKO littermate (right). **B.** Representative western blot of CNS tissues probed for NIPP1. **C.** Quantification of relative NIPP1 levels in hippocampal tissue normalized to tubulin (twotailed unpaired t-test; p = 0.0002, n = 3 animals). **D.** Quantification of relative NIPP1 levels in cortical tissue normalized to tubulin (two-tailed unpaired t-test; p < 0.0001, n = 3 animal pairs). Data are mean ± SEM.; *p<0.05, **p<0.01, ***p<0.001, ****p< 0.0001.

Since NIPP1 has been shown to influence tau processing *in vitro*, we commenced our investigation by assessing various tau phospho-epitopes. We found a drastic increase in tau phosphorylation, along with a robust change in the gel mobility shift of total tau (Fig. 2). The phosphorylation of tau at AT8 sites, Thr 181, Thr231 and Ser 214, and 324 are all robustly increased in NIPP1 cKO mice (Fig. 2). Next, looking at potential mediators of tau phosphorylation, we found that PP1 activity, determined from the level of inhibitory phosphorylation of PP1 at the C-terminus (pT320) [16], is significantly decreased in the hippocampus with a strong trend present in the cortex (Fig. 3A). AKT, MAPK and GSK3β, major kinases for tau, were also observed to have increased phosphorylation in the NIPP1 cKO brain. Whereas GSK3β activity was significantly inhibited by an increase in phosphorylation at Ser9, AKT activity measured by phosphorylation at Ser473 was significantly increased (Fig. 3B). Moreover, the increase in MAPK signaling, assayed by pERK1/2 levels, did not reach statistical significance (Fig. 3B).

**Figure 2.**
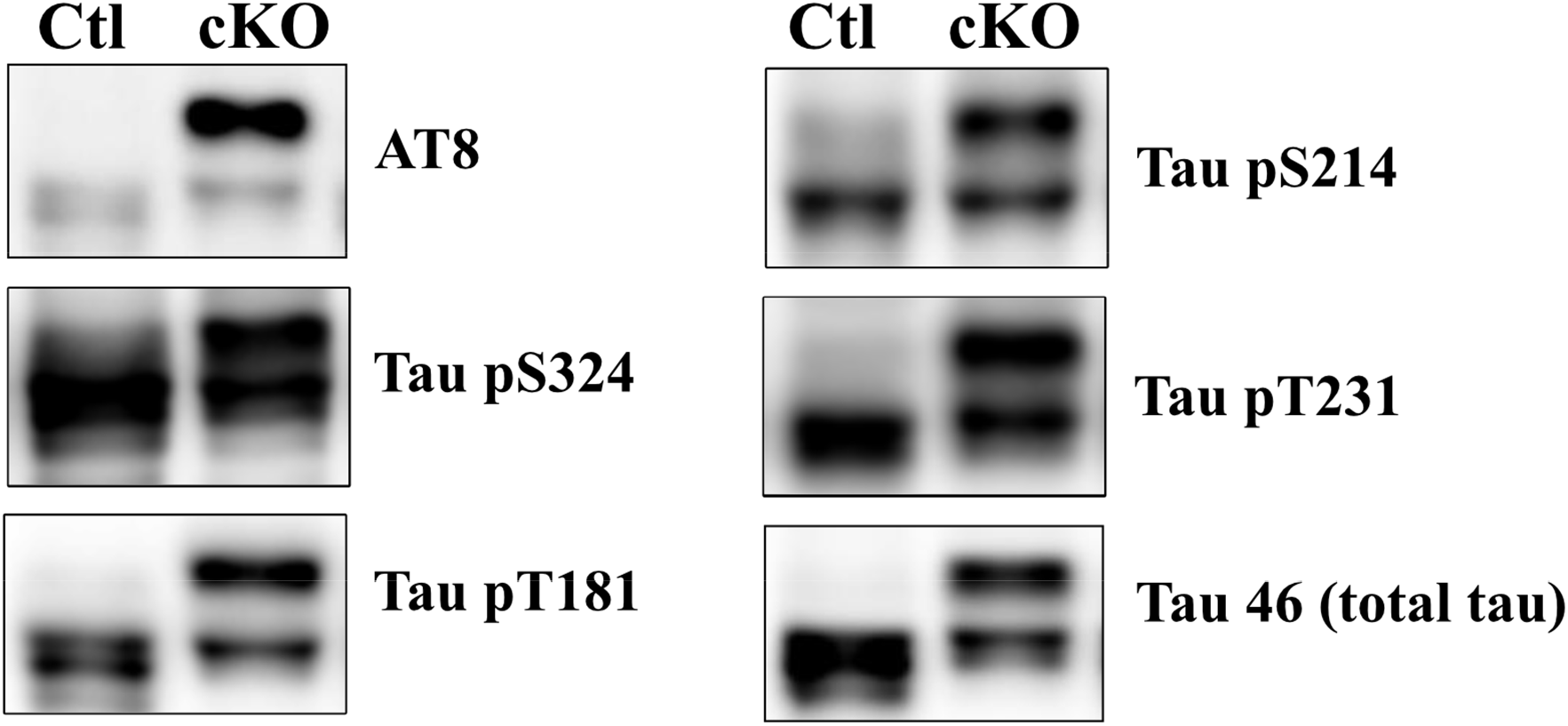
Increased tau phosphorylation in NIPP1 cKO mouse cortex. Western blot of cortical tissue from NIPP1 cKO mice probed with various antibodies specific for phospho-tau epitopes and total tau.

**Figure 3.**
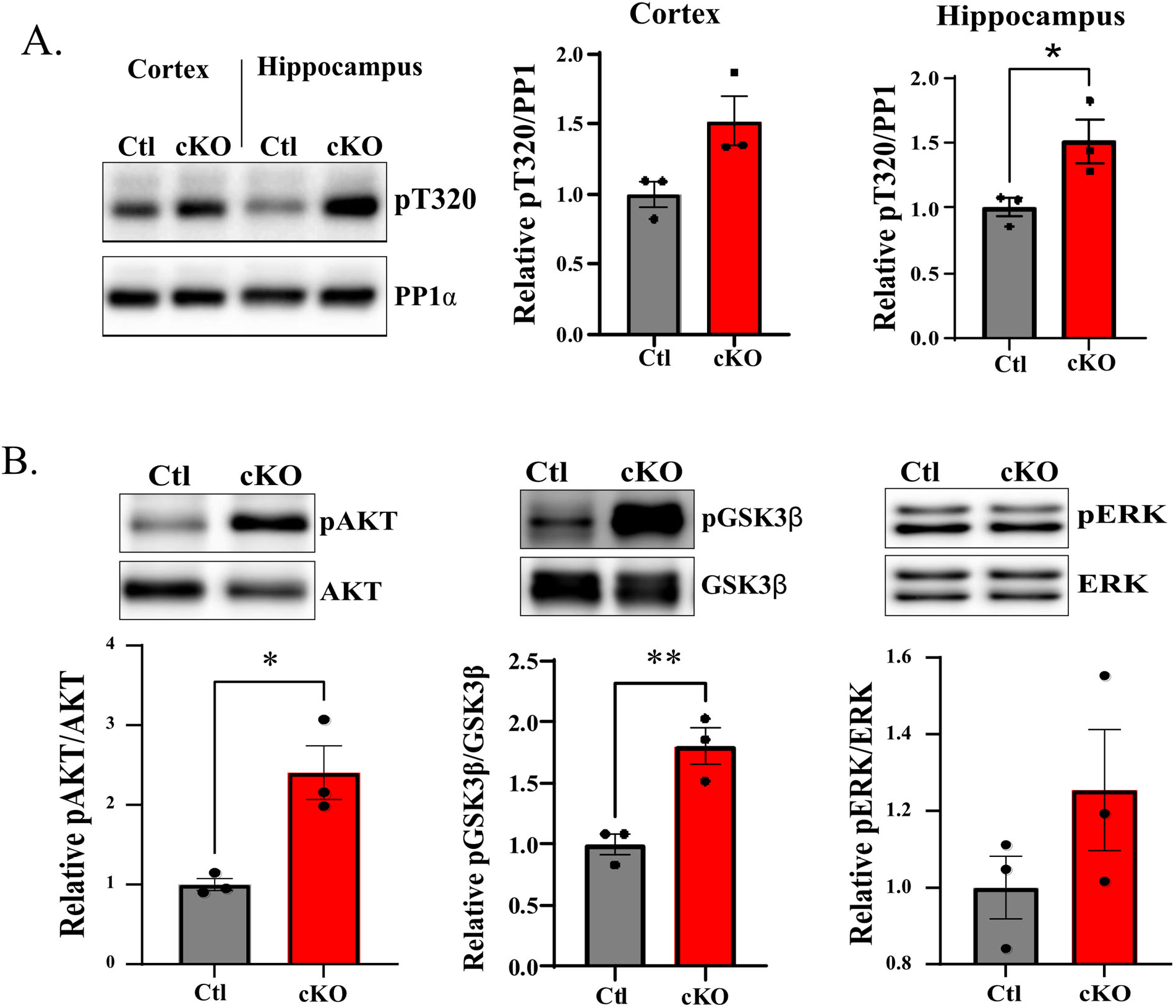
Altered activity of tau phosphatase and kinases in NIPP1 cKO mice. **A.** Western blot probed for inhibitory (pT320) PP1 epitope (left) along with quantification in cortex (Mann-Whitney test; p = 0.1) and hippocampus (two-tailed unpaired t-test; p = 0.0437, n = 3 animals) (right). **B.** Westerns blot of cortical tissue probed for various known tau kinases (left) along with quantifications (Relative pAKT; two-tailed unpaired t-test; p = 0.0153, n = 3 animals; Relative pGSK3β; two-tailed unpaired t-test; p = 0.0097, n = 3 animals; Relative pERK; two-tailed unpaired t-test; p = 0.2260, n = 3 animal pairs) (right). Data are mean ± SEM.; *p<0.05, **p<0.01.

Additionally, the NIPP1 cKO mice exhibit a severe tremor. Since abnormal myelination can cause tremor, we examined the possibility of a myelination deficit in NIPP1 cKO mice. We found that myelin basic protein (MBP) expression decreased dramatically in both HPC and cortex of NIPP1 cKO mice (Fig. 4A). One possible mechanism of myelination deficit is a change in the densities of oligodendrocyte precursor cells (OPCs) and/or mature myelinating oligodendrocytes (OLs). To assess whether these populations change in NIPP1 cKO mouse brains, we performed immunohistochemistry to measure the density of OPC-lineage cells with Olig2 and ASPA antibodies (Fig. 4BC). However, we did not observe a significant difference in mature OL or pan-OL densities within the corpus callosum denoted by ASPA, and Olig2, respectively (Fig. 4BC). In order to examine the myelin sheath directly, we performed electron microscopy (EM) on the optic nerve from NIPP1 cKO and their WT littermate controls (Fig. 5A-B). The optic nerve serves as an excellent model to study myelination due to its accessibility and its relatively high degree of myelination early in development [17]. Our data suggest no obvious difference in myelin thickness in the NIPP1 cKO mice, indicated by g-ratio measurements (Fig. 5C). Our study however, indicates a strong deficit in the percentage of myelinated axons in the NIPP1 cKO mice (Fig. 5E). We then analyzed axon caliber, a potential mechanism to explain the myelination deficit, and found a nearly identical distribution of axon calibers (data not shown). This suggests that the myelination deficit in NIPP1 cKO mouse is not driven by an increased proportion of axons that are subthreshold in size to be myelinated.

**Figure 4.**
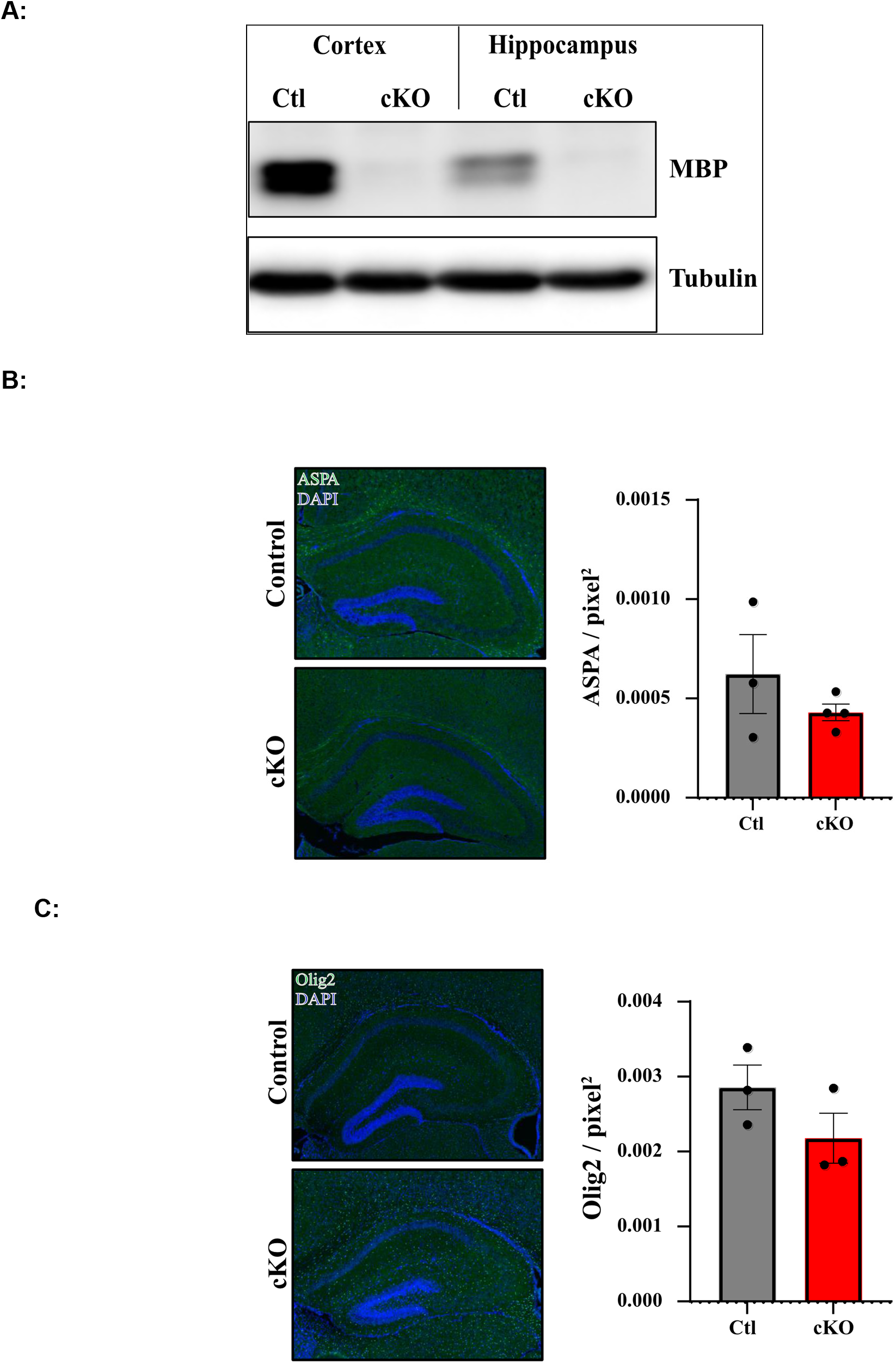
Myelin Basic Protein (MBP) level is decreased, without a change in OPC/OL densities, in NIPP1 cKO mice. **A:** Representative western blot of cortical and hippocampal tissue from NIPP1 cKO mouse with littermate control probed for MBP. (N=3 pairs); **B.** Immunostaining of brain sections from control and NIPP1 cKO mice stained with mature oligodendrocyte marker, ASPA (left). Quantification of ASPA in the corpus callosum (two-tailed unpaired t-test; p = 0.3153; ctl animals = 3, cKO animals = 4) (right). **C.** Immunostaining of brain sections from control and NIPP1 cKO mice stained with pan-oligodendrocyte marker, Olig2 (left). Quantification of Olig2 in the corpus callosum (two-tailed unpaired t-test; p = 0.2052; animal pairs = 3) (right). Data are mean ± SEM.

**Figure 5.**
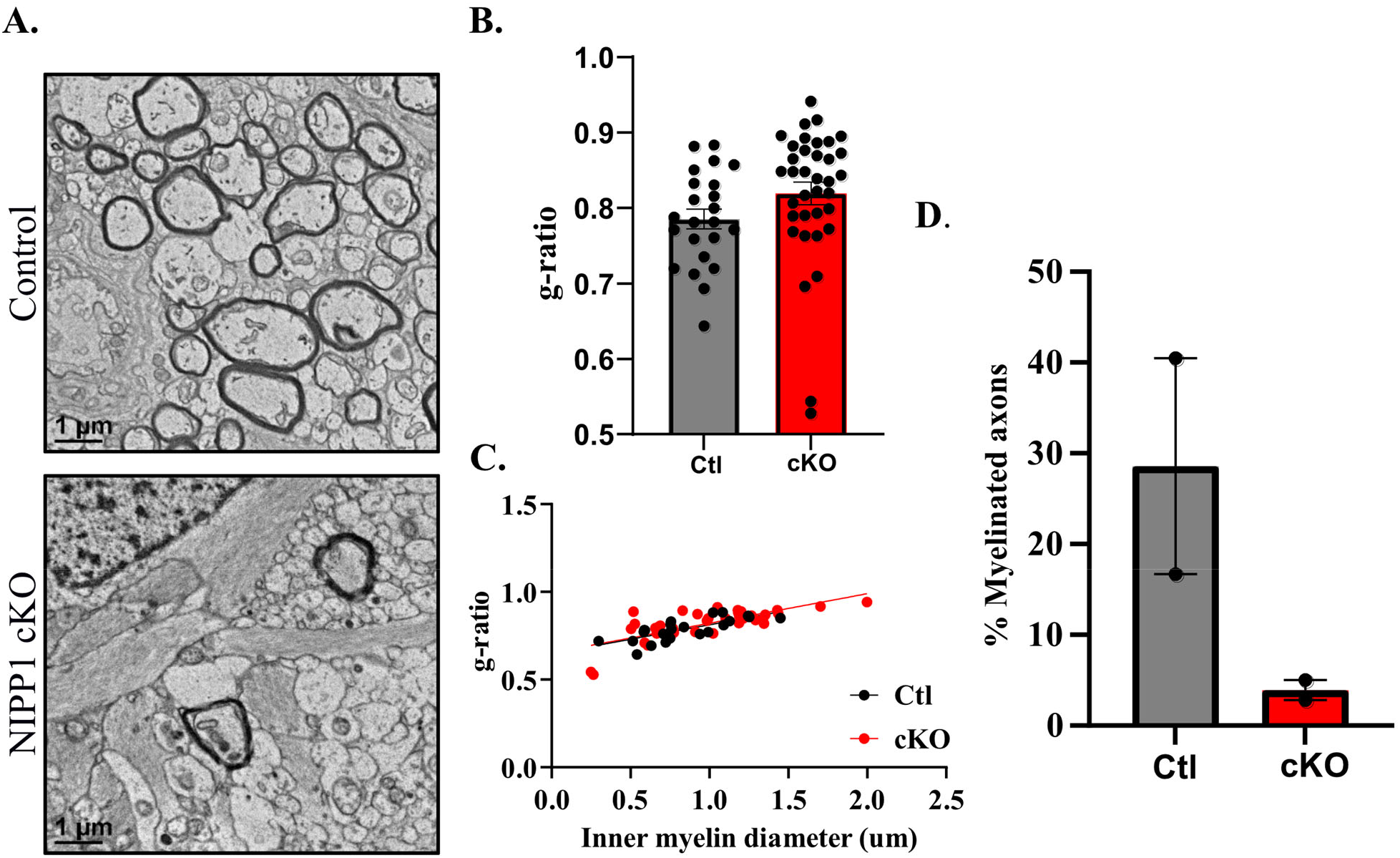
Myelin deficits in optic nerve of NIPP1 cKO mice. **A.** Representative electron micrographs of optic nerve cross sections from control and NIPP1 cKO animals respectively. **B.** mean g-ratio of myelinated axons calculated from the inner myelin diameter / outer myelin diameter (two-tailed unpaired t-test; p = 0.1168, control axons = 23; cKO axons = 36, animal pairs = 2). **C.** g-ratios plotted as a function of axon diameter, overlaid with lines of best fit. **D**. Percentage of myelinated axons. Data are mean ± SEM.

As a further test of the influence of NIPP1 on myelination, we measured compound action potentials (CAPs) from control and NIPP1 cKO mice. At ages P14-P21, the myelinated and un(pre)myelinated components of the CAPs from optic nerves are approximately comparable in amplitude. This preparation thus affords a sensitive test for myelin development in the CNS. The recordings show that deletion of NIPP1 in NPCs of the CNS significantly decreases the relative myelinated component (Fig. 6A-B). However, while conduction velocity of the myelinated component was unaffected by deletion of NIPP1, consistent with the lack of change in g-ratio, surprisingly the unmyelinated fibers conducted more slowly in the cKO mice than controls (Fig. 6C-D).

**Figure 6.**
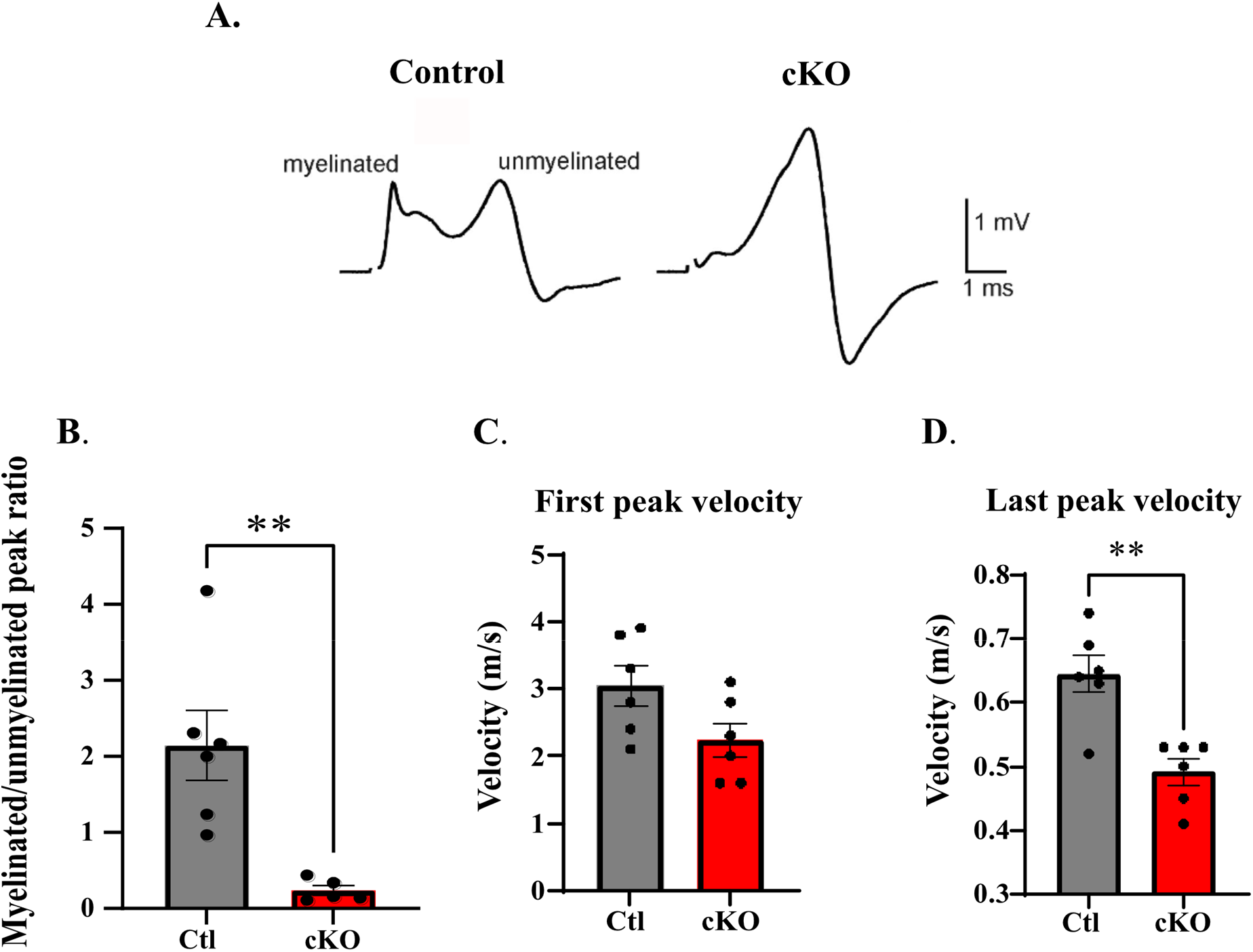
Compound action potential deficits in NIPP1 cKO mouse optic nerve. Compound action potentials from control (left) and cKO (right) optic nerves. **A**. Representative traces with stimulus artifacts removed for clarity. In each trace, the faster (first) peak represents myelinated axons, and the slower peak, unmyelinated (premyelinated) axons. **B**. Ratio of myelinated/unmyelinated peaks (two-tailed unpaired t-test; *p* = 0.0048). **C-D**. Calculated velocities of first (two-tailed unpaired t-test; *p* = 0.0653) and last peaks (two-tested unpaired t-test; *p* = 0.0018) respectively. Ctl nerves = 6, cKO nerves = 5, animal pairs = 3. Data are mean ± SEM.; *p<0.05, \*\**p*<0.01.

## Discussion

NIPP1 is a nuclear protein regulating gene expression and has been shown to play roles in many physiological and cellular processes, however, its role in CNS has not been reported yet. Homozygous deletion of the *nipp1* gene is embryonic lethal and thus necessitates the need for using a conditional knockout mouse model to study brain functions. In this study, we find deletion of NIPP1 from NPCs results in impaired development and early demise of the animal. Although a myriad of mechanisms can be involved in the animal’s failure to thrive, we began our studies investigating tau and myelin, two constituents involved in numerous neurodegenerative and neurocognitive disorders. Tau hyperphosphorylation has been shown to induce microtubule destabilization, and tau mislocalization followed by aggregation, all of which can have dire consequences for neuronal function [13]. In the present study, we report increased tau phosphorylation at AT8, S324, T181, S214 and T231 along with dysregulation of tau kinases/phosphatase activation. Decreased PP1 activation along with increased phosphorylation of AKT and GSK3β suggest a role for NIPP1 as a positive regulator of PP1. These results are reminiscent of similar effects on PP1 inhibitory phosphorylation in I-2 knockdown [16] and I-2KO studies [18]. Additionally, a gel mobility shift was observed for total tau protein. Our phospho-tau antibody blotting suggests that phosphorylation of many tau sites contribute to this shift, but we cannot exclude that there are also expression of different tau isoforms. Dysregulation of tau isoforms has been demonstrated in various tauopathies [19], and the relative levels of each isoform in NIPP1 cKO mice requires further investigation.

Another critical process that is abrogated in neurodegenerative diseases is myelination. Myelination of axons is important for the rapid transduction of action potentials, along with providing neurotrophic support to distal axons [20]. Myelin is not only critical for motor and sensory function, it also contributes to higher order cognitive functions such as learning and memory [21, 22]. Dysregulation of myelination contributes to a wide range of neuropsychiatric disorders including depression and schizophrenia [23–25]. Additionally, reduced white matter has been observed in AD patients and was reported to occur even earlier than memory deficits, suggesting the possibility that CNS myelin dysfunction could contribute to AD pathogenesis.

Although OLs have an intrinsic capability to myelinate, as demonstrated when cultured with inert nanofibers [26], this process, can be fine-tuned by specific neuron-OPC/OL and astrocyte-OPC/OL interactions. Specifically, neurons can modulate myelination by various mechanisms including: axon caliber, secretion of trophic factors and direct neuron-glial synaptic transmission [27]. Neuronal activity positively regulates CNS myelination [28–30], possibly via regulating OPC/OL densities. However, we did not observe obvious changes in OPC/OL densities in the NIPP1 cKO mice. Axon caliber also does not appear to play a major role in the myelination deficit in the NIPP1 cKO mice either. PI3K-AKT-mTOR is a prominent signaling pathway important for myelination either through its action in OPC/OLs or its action in neurons, the latter of which regulates axon caliber [17, 31]. However, we did not observe a change in mTOR activation, consistent with a no change in axon caliber in the NIPP1 cKO optic nerve (data not shown). On the contrary, the significant decrease of percentage of axon myelinated suggests the possibility of increased axon caliber for the unmyelinated axons in the NIPP1 cKO mice. However, the conduction velocity of unmyelinated axons the NIPP1 cKO mice decreased, as opposed to an increase based on a presumed increase in axon caliber. The mechanisms of failed myelination for those axons and their slowed conduction as we as the overall myelination deficit in the NIPP1 cKO mice require future investigation.

In this report, we detected tau hyperphosphorylation and CNS myelination deficits upon NIPP1 deletion in the brain. Whether tau hyperphosphorylation and CNS myelination abnormalities in NIPP1 cKO mice are related, is not known and needs to be studied in the future using cell-type specific Cre lines. Finally, mechanisms underlying decreased conduction velocity in unmyelinated component of CAPs in NIPP1 cKO also needs future investigation. In summary, we have demonstrated a critical role of NIPP1 in CNS development, and also imply a potential contribution in neurological disorders and diseases.

## Methods

### Mouse model

Nestin-Cre hemizygous mice purchased from the Jackson Laboratory were crossed with *nipp1* floxed animals generated as previously described [8]. C57BL6J mice were reared in standard housing conditions and given food/water ad libitum. All experiments were conducted at P15-21. All experiments were performed in accordance with protocols approved by University Committee on Animal Resources.

### Western blotting and antibodies

Mice were deeply anesthetized using a CO_2_ before being euthanized by decapitation with guillotine. The brain was removed quickly and different parts of the brain were rapidly dissected out on ice. The tissues were suspended in RIPA buffer and mechanically homogenized by using a polytron tissue homogenizer. Protein quantitation was performed using a BCA kit and 20 μg of total protein was loaded for SDS-PAGE as performed previously [32]. Antibodies: anti-NIPP1 (1:1000; Sigma), anti-tubulin (1:2000; Chemicon), anti-phospho-Tau (Ser202, Thr205) (AT8) (1:500, Invitrogen), anti-pTauThr181 (1:500, Cell Signaling), anti-pTauSer214 (1:500, Cell Signaling), anti-pTauThr231 (1:500, Cell Signaling), anti-Tau (46)(1:500, SCBT), anti-PP1 pT320 (1:1,000; Cell Signaling Technology), anti-PP1 (1:1,000; E-9, Santa Cruz Biotechnology, Inc.), anti-pAKT (Ser473) (1:500, Cell Signaling), anti-AKT (1:1000, Cell Signaling), anti-pGSK3β (Ser9) (1:500, Cell Signaling), anti-GSK3β (1:1000, SCBT), anti-pERK1/2 (1:500, Cell Signaling), anti-pERK1/2 (1:1000, Cell Signaling), anti-MBP (1:1000, Biorad).

### Electron microscopy

Optic nerves were immersion fixed in a combination fixative containing 2.0% paraformaldehyde/2.5% glutaraldehyde + 0.5% sucrose in 0.1M sodium cacodylate buffer (pH 7.4). After 24 hours of primary fixation the nerves were rinsed in the same buffer and post-fixed for two hours in buffered 1.0% osmium tetroxide/1.5% potassium ferrocyanide, washed in distilled water, dehydrated in a graded series of ethanol to 100% (x3), transitioned into propylene oxide, infiltrated with EPON/Araldite resin overnight, embedded into molds and polymerized for 48 hours at 60°C. The blocks were sectioned at one micron and stained with Toluidine Blue prior to thin sectioning at 70nm onto slot formvar/carbon coated nickel grids. The grids were stained with aqueous uranyl acetate and lead citrate and examined using a Hitachi 7650 TEM with an attached Gatan 11 megapixel Erlangshen digital camera and Digitalmicrograph software. Images were analyzed with *FIJI*, and G-ratios were calculated using an *ImageJ* plug-in (available at http://gratio.efil.de/), from randomly selected axons. G-ratios were calculated using the inner myelin sheath diameter (as opposed to the axon diameter, to avoid skewing data from potentially abrogated inner myelin tongues) divided by the outer myelin sheath diameter. Circumferences were manually traced.

### Electrophysiology

For complete details see [33]. Briefly, mice were euthanized with CO_2_, and optic nerves were dissected and placed in oxygenated artificial cerebral spinal fluid (ACSF) for 1 hour at room temperature. ACSF contained: 125 mM NaCl, 1.25 mM NaH_2_PO_4_, 25 mM D-glucose, 25 mM NaHCO_3_, 2.5 mM CaCl_2_, 1.3 mM MgCl_2_, and 2.5 mM KCl, and was bubbled with 95% O_2_ / 5% CO_2_. Nerves were transferred to a temperature controlled chamber, and perfused with ACSF. Each end of a nerve was drawn into a suction electrode for stimulation (at the retinal end) and recording. Stimuli were applied at 50 μsec duration, and with supramaximal current amplitudes. CAP records were low-pass filtered at 10 kHz, and fed into a data processing system for later analysis.

### Immunohistochemistry

Mice were deeply anesthetized using a combination of ketamine/xylazine. Animals were then transcardially perfused with PBS followed by 4% paraformaldehyde (PFA). Brains were carefully excised and placed in 4% PFA overnight at 4°C. Brains were then rinsed with PBS and sunk in 30% sucrose in PBS at 4°C. Brains were then carefully snap frozen in liquid nitrogen, then placed in block molds containing optimal cutting temperature (OCT) compound. Cryosections were then cut at 16μm at −17 °C. Sections were allowed to dry at room temperature overnight. Sections were then washed with PBS and placed in Liberate Antibody Binding (LAB) solution for 5 minutes. Sections were then washed 3 times with PBS and incubated in 0.5% Triton in PBS at 37 °C for 30 minutes. Slides were then washed 3 more times with PBS. 5% donkey serum in PBS was then used to block the sections for 1 hour. Primary antibodies, ASPA (1:100 Genetex 113389) or Olig-2 (1:100 Millipore AB9610) were added, and slides were incubated at 37 °C for 1 hour, then held overnight at 4 °C. Slides were then washed with PBS, and incubated for 1 hour at room temperature with Alexa 488 donkey anti-rabbit antibody (1:400, in 5% donkey serum). DAPI was used to label nuclei. Thoroughly washed slides were mounted and allowed to dry overnight before imaging on Keyence BZ-X800 fluorescent microscope. Images were randomized using a plug-in for *ImageJ* (*https://imagej.nih.gov/ij/macros/Filename_Randomizer.txt*) and the corpus callosum of each image was carefully traced by hand. Only cells within that region of interest were then counted

### Statistical analyses

The Shapiro-Wilk normality test was applied to verify normality of data. Unpaired two-tailed t-tests with alpha set to 0.05 were applied to parametric data to compare the differences between two groups. The Mann-Whitney U test was applied for sets where both groups rejected the null hypothesis of the Shapiro-Wilk test. Averages were denoted as mean ± SEM in all bar graphs. All statistical tests were performed using *GraphPad Prism9.3* software.

## Acknowledgements

This work is supported by the National Institutes of Health (NIH) (R01 MH109719) and National Science Foundation (NSF) (IOS-1457336) to HX; and the NIH (F30 MH122046) to KF. We thank the technical help of Karen Bentley, director of Electron Microscopy Core of URMC and Drs. Mathieu Bollen and Aleyde Van Eynde for providing the floxed NIPP1 mice.

